# Uncertainty-aware single-cell annotation with a hierarchical reject option

**DOI:** 10.1101/2023.09.25.559294

**Authors:** Lauren Theunissen, Thomas Mortier, Yvan Saeys, Willem Waegeman

## Abstract

Automatic cell type annotation methods assign cell type labels to new datasets by extracting relationships from a reference RNA-seq dataset. However, due to the limited resolution of gene expression features, there is always uncertainty present in the label assignment. To enhance the reliability and robustness of annotation, most machine learning methods address this uncertainty by providing a full reject option, i.e. when the predicted confidence score of a cell type label falls below a user-defined threshold, no label is assigned and no prediction is made. As a better alternative, some methods deploy hierarchical models and consider a so-called partial rejection by returning internal nodes of the hierarchy as label assignment. However, because a detailed experimental analysis of various rejection approaches is missing in the literature, there is no consensus on best practices, superiority of certain methods, and potential drawbacks associated with rejection. We evaluate three annotation approaches (1) full rejection (2) partial rejection and (3) no rejection for both flat and hierarchical probabilistic classifiers. Our findings indicate that hierarchical classifiers are superior when rejection is applied, with partial rejection being the preferred rejection approach, as it preserves a significant amount of label information. For optimal rejection implementation, the rejection threshold should be determined through careful examination of a method’s rejection behavior. Without rejection, flat and hierarchical annotation perform equally well, as long as the cell type hierarchy accurately captures transcriptomic relationships.

## 1 Introduction

Due to the structural and functional heterogeneity of individual cells, cell categorization into cell types is a crucial pre-processing step before further downstream analysis of single-cell RNA-seq data [Luecken and Theis, 2019]. Cell types represent cell groups that distinctly exhibit similar functionality and structure. Their identification enables biological interpretation of the data and provides guidance for further analysis of the transcriptomics data [Heumos et al., 2023, Clarke et al., 2021, Hongkui Zeng, 2022].

There are two different ways to annotate single-cell experiments with cell type information: (1) the cells of an experiment can be clustered in an unsupervised manner and subsequently manually annotated by experts with the help of known marker gene lists and domain knowledge, or (2) the cells can be annotated in an automated fashion, with supervised learning algorithms that rely on marker-genes lists or reference cell datasets. Due to the subjectivity, irreproducibility and time-consuming nature of manual annotation pipelines, automated methods are increasingly used [Clarke et al., 2021, Pasquini et al., 2021].

However, the drawback of automated annotation methods is that they are not good at handling annotation uncertainty (a.k.a. predictive uncertainty in machine learning). This uncertainty can be caused by multiple factors: incomplete reference datasets, misalignment with experimental conditions, incomplete marker gene lists, imbalance in the presence of cell populations, limited information-content of the datasets’ features etc. On top of this, many cell types do not have well-characterized gene expression signatures, which already makes them hard to annotate [Clarke et al., 2021]. Ideally, the uncertainty resulting from these factors would be accounted for in the automatic labeling process, so that the final label assignment can be perceived as trustworthy. Some automatic annotation methods handle this uncertainty by implementing a reject option during labeling. If the labeling uncertainty becomes too large an unknown label is assigned instead of a cell type label. The majority of methods that have a reject option base their rejection process on thresholding a prediction confidence measure [Li, Yingxin et al., 2020, Michielsen et al., 2021, Pliner et al., 2019, de Kanter et al., 2019, Kiselev et al., 2018, Alquicira-Hernandez et al., 2019, Cao et al., 2020].

Furthermore, the relationship between cell types is often hierarchical in nature [Wu and Wu, 2020]. Therefore, several automatic annotation tools exist that utilize these hierarchical relationships to more accurately annotate single-cell datasets. Hierarchical classifiers annotate sequenced cells with increasing level of detail by traversing the hierarchy, making annotation decisions in a step-wise manner. Both marker-based and supervised hierarchical annotation tools exist. Some of these tools have the ability to self-construct cell type hierarchies [de Kanter et al., 2019, Li, Yingxin et al., 2020, Bassel Ghadar and Subhajyoti De, 2022], while others require that the end user provides a hierarchy [Li, Yingxin et al., 2020, Kaymaz et al., 2021, Pliner et al., 2019, Galdos et al., 2022, Prummer et al., 2022]. One can also use existing cell type hierarchies, such as the cell ontology [Bernstein et al., 2021, Wang et al., 2021].

Annotation methods that incorporate cell type hierarchies are appealing, because they have the ability to implement a “partial” reject option [de Kanter et al., 2019, Pliner et al., 2019, Michielsen et al., 2021, Bi and Kwok, 2015]. They can reject cell types labels inside the hierarchy, which allows them to assign intermediate cell type labels present higher up the hierarchy if the uncertainty of these intermediate labels is not too high. They thus preform a partial reject. In this way, these methods are able to retain more label information compared to non-hierarchical or flat models [Wang and Casasent, 2009, Sun and Lim, 2001, Ceci and Malerba, 2007].

Although a partial reject option has been introduced in a few hierarchical classifiers for single-cell annotation, an in-depth empirical evaluation is currently lacking in the literature. This paper intends to bridge this gap by evaluation three strategies for cell annotation with flat and hierarchical classifiers: 1) label assignment without rejection 2) label assignment with full rejection and 3) label assignment with partial rejection. Label assignment with full rejection is the most common rejection method in the literature and will result in the assignment of an ’unknown’ label in the presence of uncertainty. Label assignment with partial rejection is implemented with the help of a cell type hierarchy and can lead to a more general label assignment, that is present at an intermediate level in the hierarchy. We investigate the impact of these rejection approaches on the annotation task and formulate best-practices for rejection implementation. We introduce accuracy-rejection curves as a handy tool for the practical determination of the optimal rejection threshold and to compare rejection behavior across models. We also compare flat and hierarchical annotation performance without rejection and look at the interpretability of these models.

## 2 Methods

### 2.1 Data sets and preprocessing

We analyzed five single-cell transcriptomics datasets with publicly available cell hierarchies: the AMB dataset [Tasic et al., 2018], the COVID dataset [Chan Zuckerberg Initiative Single-Cell COVID-19 Consortia et al., 2020], the AzimuthPBMC dataset [Stuart et al., 2019], the Flyhead dataset and the Flybody dataset [Li et al., 2022]. These datasets differ in size, species, type of hierarchy and complexity. The AMB and COVID dataset were labeled by unsupervised clustering and marker gene inspection. The Azimuth PBMC dataset was generated by Stuart et al. [2019], integrating an RNA-seq and CITE-seq dataset of human PBMC. In our analyses, only the RNA-seq information was used. The Flyhead and Flybody dataset were manually annotated by experts from 40 laboratories through consensus-voting according to the Flybase ontology conventions [Li et al., 2022, Gramates et al., 2022]. - Characteristics of the datasets can be found in Table S1. For all the datasets, cell type populations with less than 10 observations were filtered out, together with doublets, unknown cells, and non-uniquely specified cells. The label hierarchies of all the datasets can be found back in Figures S18 - S22. For the datasets with ontology-based annotation, the annotation was shortened if it did not affect other labels, e.g. if one label had 10 unique extra levels and thus labels. These labels were deleted and subsequently the full label became shorter. Prior to all analyses the count data was log-transformed with the following function: log_2_(*x* + 1).

### 2.2 Formal problem definition and flat annotation

In machine learning terms, supervised cell type annotation translates to a multiclass classification problem, where the goal is to assign one cell type label to each individually sequenced cell. Formally speaking, we consider a labeled training dataset *𝒟* = (**x**_1_*, y*_1_), (**x**_2_*, y*_2_), …, (**x***_n_, y_n_*), consisting of *n* couples, with **x***_i_* a *p*-dimensional feature vector that represents gene expression for *p* genes, and *y_i_* the corresponding label. In the realm of uncertainty, the ground-truth label *y* that corresponds to a feature vector **x** cannot be determined in a deterministic manner. Hence, we assume that the dataset *𝒟* is i.i.d. sampled according to an underlying joint distribution *P* over *X × Y* with *X* the feature space and *Y* = *{c*_1_*, …, c_K_}* the label space, containing *K* unique classes.

Discriminative probabilistic classifiers express uncertainty in the label *c* that corresponds to a feature vector **x** by estimating *P* (*y* = *c |* **x**) during model training. In our experiments we used three classical classifiers of that kind: logistic regression (LR), the linear support vector machine (SVM) and random forests (RF). Standard implementations of these classifiers in scikit-learn [Pedregosa et al., 2011] were used – see supplementary material for implementation details. Logistic regression inherently learns to estimate *P* (*y* = *c |* **x**) during model training by optimizing the log-loss. For the linear SVM model, probabilities can be obtained through a softmax operation on the decision scores produced by the standard one-versus-all decomposition. For the random forest model, the class probability in a single tree is estimated as the fraction of samples of that class in the leaf assigned to the feature vector. Subsequently, the class probabilities for the entire random forest are computed as the mean class probabilities across the trees present in the forest.

In the remainder of this paper, these three classical methods will be referred to as ’flat’ classifiers, because they do not take hierarchical information into account to estimate the class probabilities. Moreover, these methods estimate *P* (*y* = *c |* **x**) using a single probabilistic classifier. After training, labels can be assigned to a new instance **x** by simply choosing the class with the highest probability:

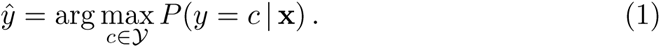

### 2.3 Hierarchical annotation

#### 2.3.1 Model training

Unlike flat classifiers, hierarchical classifiers take relationships between labels into account by exploiting a predefined label hierarchy under the form of a tree structure *T* that contains *M* nodes *V_T_* = *{v*_1_*, …, v_M_ }*. Each node represents a set of classes, and nodes near the bottom of the tree structure represent labels with a more fine-grained granularity. As special cases, the root node *v*_1_ represents the entire class space *Y*, whereas the leave nodes coincide with the individual classes. In the context of single-cell transcriptomics, each node represents a cell type label – see Figure 1.A for an intuitive example.

**Figure 1:**
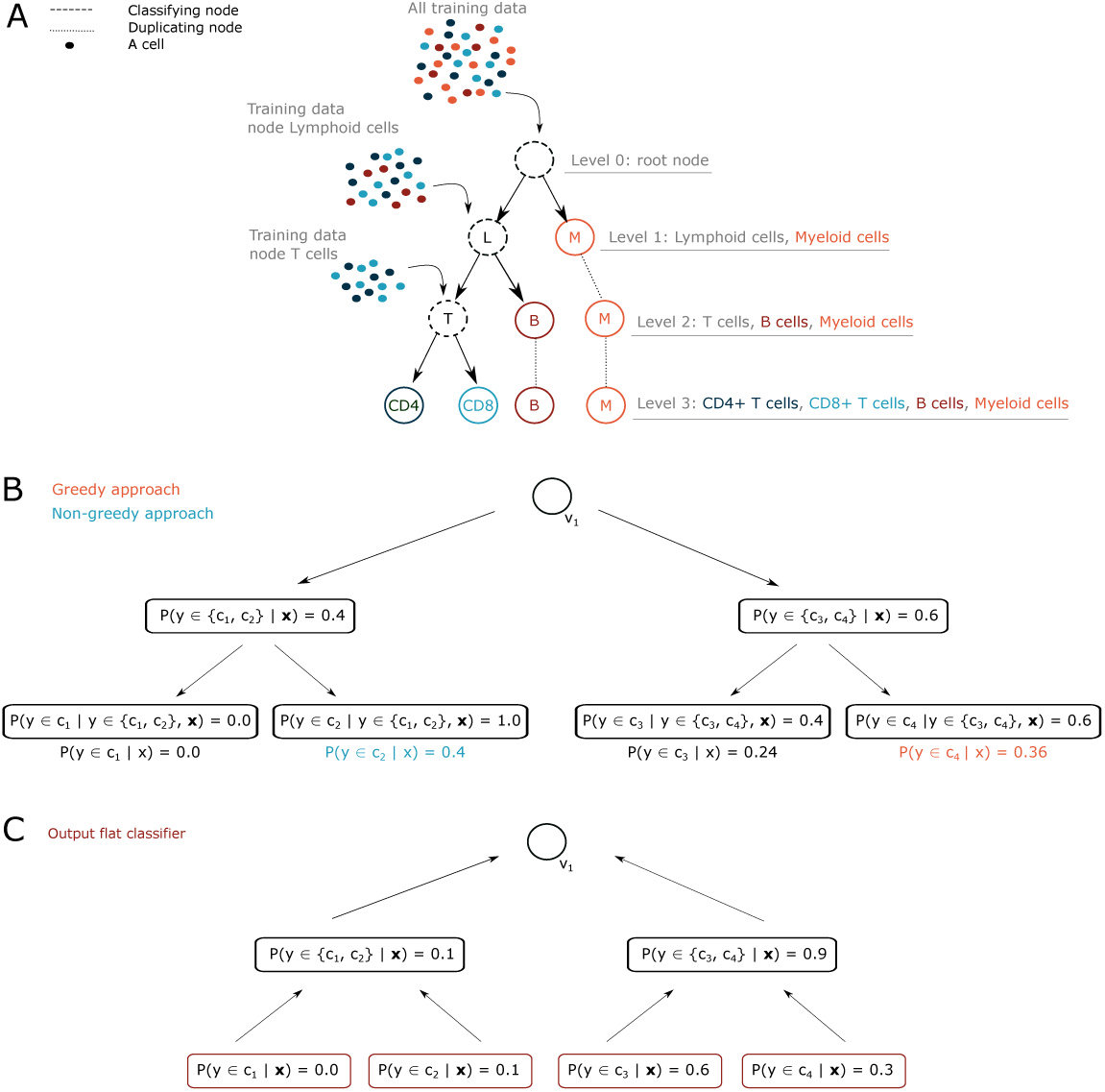
A. Schematic overview of the training of a hierarchical classifier and the division of the training data during this task. B. Example illustrating the difference between greedy versus non-greedy label assignment with hierarchical annotation. With greedy label assignment, only the path with the highest probability scores in the hierarchy is followed. With non-greedy label assignment, all possible prediction paths are traversed and only the end score is considered for the final label assignment. C. Illustration of how intermediate node probabilities for a cell type hierarchy are reconstructed for bottom-up label assignment. The intermediate node probabilities are calculated by summing all the probabilities of the children nodes. This information can then be used to perform label rejection along the hierarchy and to construct accuracy-rejection curves.

Probabilistic hierarchical classifiers learn *P* (*y* = *c |* **x**) in a hierarchical manner [Bi and Kwok, 2015]. The probability mass *P* (*y ∈ v |* **x**) for an instance **x** assigned to node *v* can be computed using the chain rule of probability:

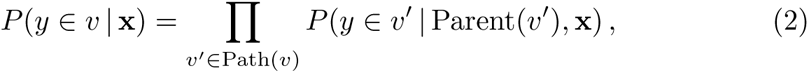

with Parent(*v*) the parent of node *v*, and Path(*v*) the set of nodes on the path connecting the node *v* and the root of the tree structure. For the root node *v*_1_ we have *P* (*y ∈ v*_1_ *|* Parent(*v*_1_), **x**) = 1. In each node of the hierarchy, one can train any standard multi-class probabilistic classifier. In the machine learning literature, methods of that kind are known with different names, such as nested dichotomies [Frank and Kramer, 2004, Melnikov and Hüllermeier, 2018], conditional probability estimation trees [Beygelzimer et al., 2009] and probabilistic classifier trees [Dembczyński et al., 2016]. The notion *local classifier per parent node* is also used, but then usually no restriction is made to probabilistic models [Silla and Freitas, 2011].

Training typically starts at the root node, where a probabilistic classifier is trained on all training samples to distinguish labels that are one level lower in the hierarchy. For the example in Figure 1.A, this corresponds to the distinction of the orange samples (Myeloid cells) from the other samples. Then, all the training samples are split according to their ground truth level 1 label and passed to the corresponding label node in the hierarchy, where a new classifier will be constructed that learns to distinguish between the labels one level lower, as illustrated in Figure 1.A. The training process ends when a probabilistic classifier has been fitted in every internal node of the class hierarchy. In our experiments, logistic regression, random forests and linear SVM models were implemented as independent classifiers inside the hierarchy. Within a single hierarchical model the same type of classifier was considered for all the internal nodes of the hierarchy.

#### 2.3.2 Greedy and non-greedy label assignment

After training, the model can be used for label assignment of (new) test instances. Similar to Eqn. (1) for flat classifiers, we aim to return for an instance the class with highest probability. However, computing all conditional class probabilites *P* (*y* = *c |* **x**) is time comsuming, because the chain rule of probability in Eqn. (2) needs to be invoked for every class *c*. Therefore, in the literature, several algorithms have been proposed to compute Eqn. (1) more efficiently for hierarchical classifiers. In our experiments, we implemented two algorithms, further referred to as greedy and non-greedy label assignment. The two approaches were originally proposed in [Read et al., 2009] and [Dembczyński et al., 2012], respectively, for multi-label classification. Some years later they were extended for multi-class classification [Bi and Kwok, 2015, Dembczyński et al., 2016, Mortier et al., 2021]. Pseudocode of the two algorithms can be found in the Algorithms S1 and S2 in the supplementary information.

The main difference between greedy and non-greedy class assignment is illustrated in Figure 1.B. In the greedy approach only one path in the hierarchy is traversed. At the start, samples are passed to the classifier present in the root node. Then, in a “greedy” way, the child with the highest probability is chosen, and the sample is passed to this child. This process is repeated until a leaf node is reached. The second approach is the non-greedy or Bayes-optimal approach, where multiple paths starting from the root are explored, with a best-first search algorithm. In this algorithm, a priority queue orders candidate nodes for exploration according to their probability *P* (*y ∈ v |* **x**). In every step of the algorithm, the most promising node is visited, and the path to that node is further explored using Eqn. (2). The algorithm ends when the first leaf is found. One can guarantee that this leaf corresponds to the solution of Eqn. (1), i.e., the leaf node with highest probability among all leaves.

The greedy and non-greedy methods might return different predictions. For the example in Figure 1.B, leaf *c*_4_ is found using the greedy approach, whereas leaf *c*_2_ is obtained using the non-greedy approach. From a theoretical perspective, the non-greedy approach will always find the Bayes-optimal solution of Eqn. (1), at the cost of a longer runtime, while the greedy approach might yield worse predictions in a faster way. We refer to Dembczyński et al. [2012] for more detailed theoretical results on the predictive performance and runtime of both methods.

### 2.4 Full and partial reject option

Without hierarchical information, standard multi-class classifiers can only consider a standard reject option. If the uncertainty for a test instance **x** is too high, i.e. when the highest class probability *P* (*y* = *c |* **x**) over all classes *c* drops below a predefined threshold, no class label is assigned, and the instance is classified as “unknown”. We further refer to this approach as the “full reject” option. A full reject option can be implemented based on the class probabilities calculated by flat annotation or hierarchical annotation with greedy or non-greedy label assignment.

As discussed in the introduction, class hierarchies make it possible to consider a “partial reject” option. Then, an instance can be classified into an internal node of the hierarchy, when the probability of belonging to one of the children of that node drops below a predefined threshold. A partial reject option can be implemented in two ways: a first way in which the hierarchy is traversed top-down, using hierarchical classifiers, and a second way in which the hierarchy is traversed bottom-up, using flat classifiers.

The top-down approach can be combined with greedy (Algorithm S1) and non-greedy label assignment (Algorithm S2). For greedy label assignment, annotation along the hierarchy is stopped if the overall confidence score of the label becomes lower than the rejection threshold *t_r_* (see Algorithm S1). For the non-greedy approach, described in Algorithm S2), all possible label paths are restricted to those paths that do not violate the rejection threshold *t_r_*. With a best-first search algorithm, exploration of all label paths continues until the path probability drops below the rejection threshold. Similar to the Bayes-optimal algorithm without rejection (i.e. Algorithm S2 with *t_r_*= 0), one can guarantee that the path with the highest path probability that does not violate the rejection threshold is returned [Mortier et al., 2021].

The bottom-up approach is used for flat annotation models and illustrated in Algorithm S3. Initially, annotation is performed at the lowest level of the hierarchy, because the conditional class probabilities *P* (*y* = *c |* **x**) can be retrieved from the flat classifier. The algorithm picks the leaf *c* for which *P* (*y* = *c |* **x**) is the highest. If that probability is lower than the predefined threshold *t_r_*, the algorithm moves to the parent *v* of that node. Similar to the top-down approach, the probability of that parent is the sum of its children:

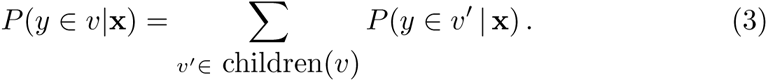

If the probability of that parent is high enough, the algorithm stops. Otherwise, the parent of that parent is visited. In the worst case, the algorithm stops in the root node, which corresponds to a full reject, as defined above.

To inspect the rejection behavior of the flat and hierarchical models, accuracy-rejection curves [Nadeem et al., 2010] are constructed for each optimized model. To construct these curves, the probability scores *P* (*y ∈ v*_1_*|***x**), *P* (*y ∈ v*_2_*|v*_1_, **x**), …, *P* (*y ∈ v_q_|v*_1_*…v_q−_*_1_, **x**) of all the labels in the nodes that make up the final predictions need to be recorded. The accuracy-rejection curves are calculated by varying the rejection threshold *t_r_* between 0 and 1.0 with a stepsize of 0.1 and performing label assignment with rejection. With these results, the accuracy score is then calculated based on the cells whose labels were not rejected.

### 2.5 Model implementation

In general, all the models were evaluated with 5-fold cross-validation (80% training and 20% test). Per fold, the training set was then further split into a training and validation set (75%/25%). The validation set was used to evaluate regularization hyperparameters of the machine learning models and feature selection hyperparameters. The test set was used to give unbiased estimates of the model performance. For the hierarchical classifiers, one set of hyperparameters was optimized for every classifier in the hierarchy, due to the computational burden.

As a result, all trained hierarchical models contained the same hyperparameter values in their nodes.

Depending on the subject of evaluation during this study, different evaluation metrics were used within the cross-validation scheme to pick the best hyperparameter combination. During rejection evaluation (Section 2.4), the log-loss was used as scoring metric. During model performance without rejection (Section 3.3) and interpretability (Section 3.5) evaluation, the accuracy score was used.

## 3 Results

### 3.1 Hierarchical annotation rejects less labels than flat annotation

Full label rejection is evaluated for flat and hierarchical annotation with the help of accuracy-rejection curves that are depicted in Figure 2 [Nadeem et al., 2010]. These curves are constructed for flat and non-greedy hierarchical annotation with three classifiers: the logistic regression classifier, the random forests classifier and the linear SVM classifier, across the 5 datasets introduced in Section 2.1. Figure 2 visualizes, besides the accuracy score across the varying rejection thresholds, also the rejection percentage for every rejection threshold. Figure 2 shows that the rejection behavior of the three classifiers, irrespective of the annotation type, varies substantially. The logistic regression classifier exhibits substantially fewer rejections at lower rejection thresholds than the linear SVM and random forests classifier. This results in a relatively flat accuracy-rejection curve with a sharp incline near the higher end of the threshold range, suggesting that the logistic regression classifier tends to be more confident in its predictions than the other classifiers. On the contrary, the random forests and linear SVM classifier display a more gradual rejection curve, indicating more variability in their confidence scores. This could indicate that the logistic regression classifier is overconfident. However, the accuracy scores obtained with the logistic regression classifier across the 5 datasets for the different rejection thresholds are quite high (80-90%). So the high confidence scores of the logistic regression are accompanied by a lot of correct predictions, which debunks the hypothesis of overconfidence. Overall, the difference in behavior has primarily practical implications, as the optimal rejection threshold across these three classifiers will differ quite severely. These results indicate the importance of constructing these accuracy-rejection curves for every specific use case prior to rejection implementation.

**Figure 2:**
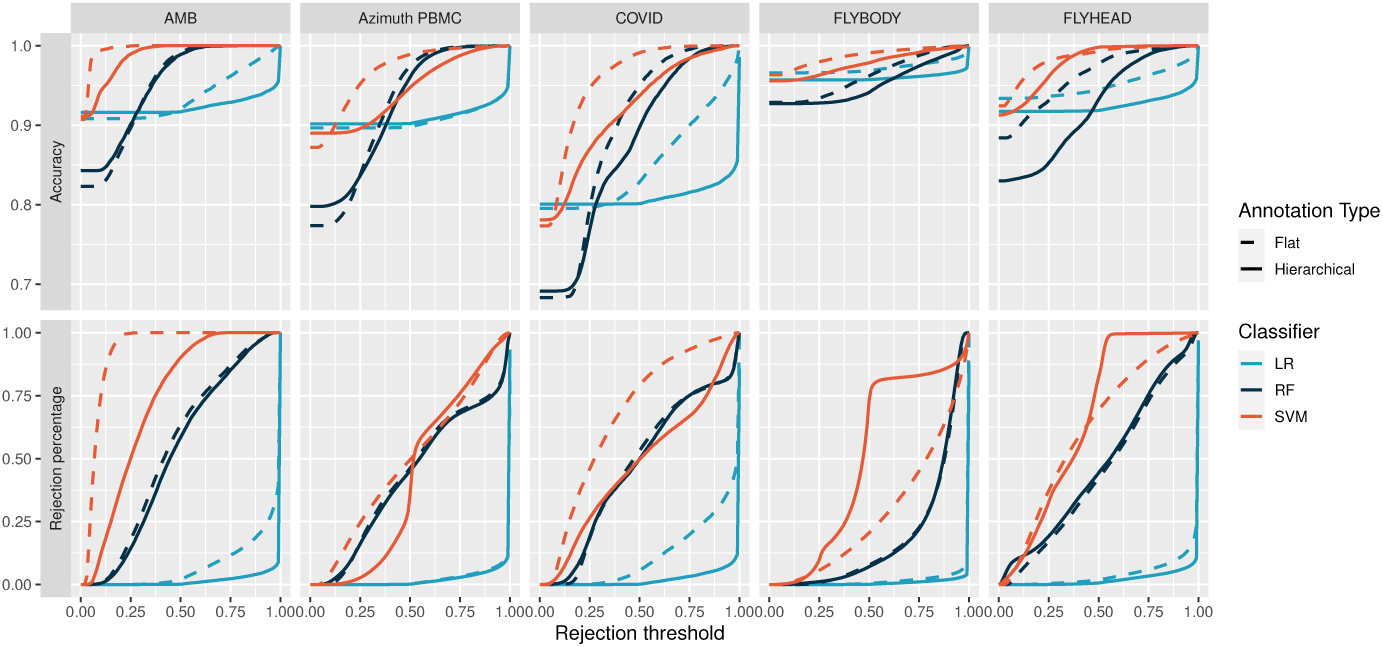
Accuracy-rejection curves of flat and hierarchical annotation with non-greedy prediction of the AMB, Azimuth PBMC, COVID, Flybody and Flyhead datasets with logistic regression (LR), random forests (RF) and linear SVM (SVM) classifiers. The curves in the top row represent the accuracy score of the non-rejected label given a rejection threshold value. The curves on the bottom row represent the percentage of rejected labels for rejection threshold value.

Comparison of the behavior of flat and hierarchical annotation across the datasets, given the same classifiers, shows that flat annotation tends to reject more at lower rejection thresholds than hierarchical annotation. One exception to this is the linear SVM classifier applied to the Flybody and Flyhead datasets. These results indicate that flat annotation, despite achieving similar performance, preserves less information than hierarchical annotation. Therefore, hierarchical annotation appears to be the better annotation approach when full rejection is implemented.

Figure S4 visualizes accuracy-rejection curves for greedy and non-greedy hierarchical annotation and shows that the curves for these two approaches almost completely coincide. So the conclusions formulated above hold true for hierarchical annotation in general, irrespective of the label assignment approach.

In practice, accuracy-rejection curves, such as Figure 2 and Figure S4, hold substantial practical value. The introduction of a reject option during the annotation process inherently introduces a trade-off between label certainty and label assignment. With these curves, the users gain a clear image of this tradeoff and can easily decide on an optimal decision for their problem at hand. As a result, we strongly recommend to always generate these curves with single-cell annotation methods that implement a rejection approach, so that an informed choice regarding the rejection threshold can be made.

### 3.2 Partial rejection retains substantially more label information than full rejection

To compare full and partial rejection for flat and non-greedy hierarchical annotation, we visualized in Figure 3 for the same datasets and classifiers as the previous section the difference in the percentage of ’unknown’ labels after rejection at a given rejection threshold. In Figure 3, the solid red line indicates the results of non-greedy hierarchical annotation with full rejection, the dashed red line the results of non-greedy top-down label assignment. The solid and dashed blue lines indicate respectively the results of flat annotation with full rejection and partial rejection (i.e. bottom-up label assignment). Note that the full rejection results (the red lines) in Figure 3 are equivalent to the results visualized in the bottom row of Figure 2, as the percentage of rejection is equal to the percentage of unknown labels.

**Figure 3:**
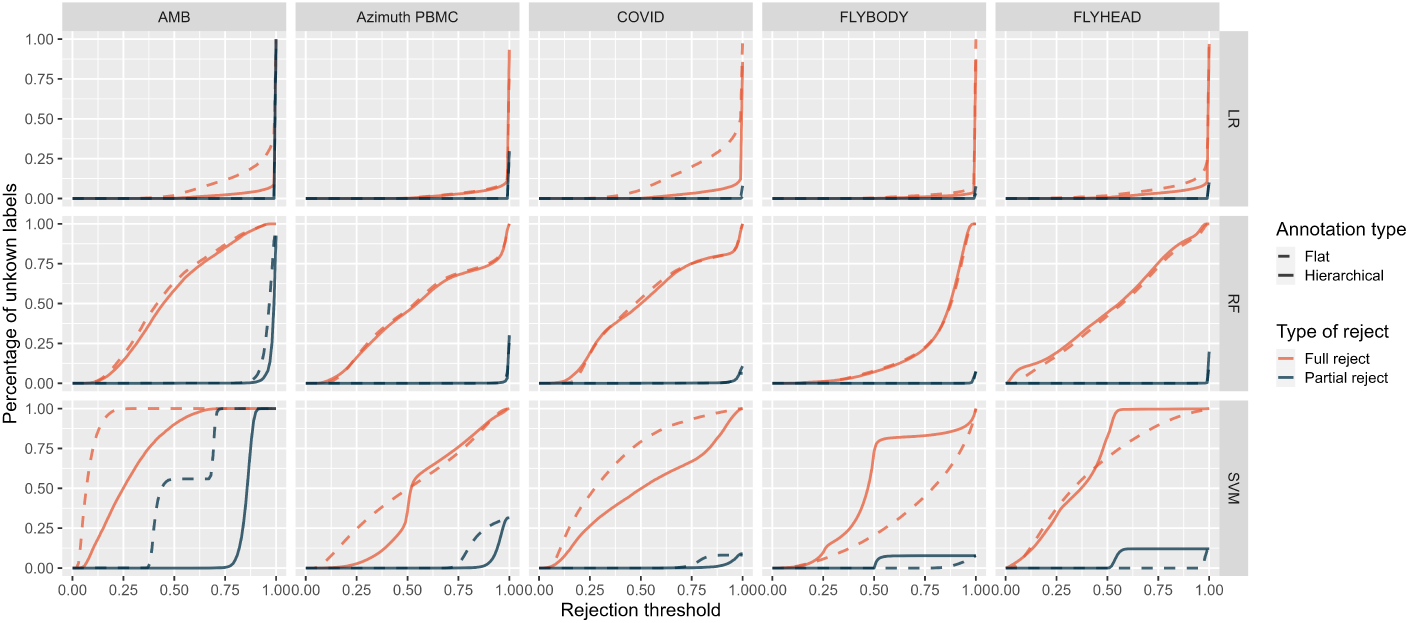
A comparison of the percentage of unknown labels assigned during the annotation task when partial and full rejection is implemented during non-greedy hierarchical and flat annotation. Evaluation is performed with three classifiers (logistic regression, random forests and linear SVM) across 5 datasets (AMB, Azimuth PBMC, COVID, Flyhead and Flybody).

Figure 3 shows that there is a substantial increase in the amount of label information that is retained with partial rejection, both with non-greedy top-down and bottom-up label assignment, in comparison to full rejection. The same comparison as Figure 3 is made for greedy and non-greedy top-down label assignment in Figure S5 and indicates no difference in the rejection behavior of the greedy and non-greedy approach for top-down label assignment.

A deeper examination of the label content after partial rejection with the help of Figures S6 and S7, shows more pronounced differences between the classifiers and annotation methods. The logistic regression classifier assigns very few intermediate labels. The linear SVM and random forests classifier assign substantially more intermediate labels and showcase a distinct stepping-like pattern: the higher the rejection threshold, the more severely labels are rejected. Comparison of flat and hierarchical annotation, across the datasets and classifiers based on Figures S6 and S7, learns that flat annotation tends to reject more severely (a.k.a to higher levels in the hierarchy) than hierarchical annotation, resulting in less specific annotation for a given threshold.

Overall, these results augment the two conclusions made in the previous section. (1) There is substantial variability present in the rejection behavior between the different classifiers and this highlights the necessity for the inspection of rejection behavior for a specific use-case prioir to rejection implementation (2) Hierarchical annotation tends to reject less and less severely than flat annotation in the case of uncertainty, regardless of the annotation type. The results also indicate the ability of partial rejection to retain substantially more label information in the case of uncertainty, regardless of the annotation type.

### 3.3 Flat and hierarchical annotation without rejection perform similarly well, given a good cell type hierarchy

We evaluated the prediction performance of the classifiers under the two annotation types, flat and hierarchical, without taking the rejection process into account on the 5 datasets that were also used in the previous sections. The prediction performance of the flat and hierarchical annotation models are compared with the average accuracy score across the 5-fold cross-validation and the overall balanced accuracy score, i.e. the average of recall over each cell type population. The results of this comparison for flat and greedy hierarchical annotation can be found in Figure S8.

Generally, for the manually annotated datasets, the AMB, Azimuth PBMC and COVID dataset, the overall performance of hierarchical annotation matches that of flat annotation both in terms of the balanced accuracy score (Fig. S8.A) and the accuracy score (Fig. S9). The balanced accuracy score is important to take into account due to the large class imbalance that is inherently present in all the single-cell datasets. This imbalance can create a bias in the accuracy score as the complete misclassification of smaller classes will not be reflected. For the datasets with ontology-base annotation, the performance of hierarchical annotation drops.

The impact of feature selection on both annotation strategies is also evaluated with the implementation of the F-test as feature selection method and highly variable gene (HVG) selection [Satija et al., 2015] – more details can be found in Figure S9. The results (Fig. S9) indicate that feature selection consistently improves annotation performance. Both the F-test and HVG selection seem to outperform each other very slightly depending on the dataset and classifier, making both methods equally worthwhile to incorporate in future cell type annotations.

A comparison of greedy and non-greedy hierarchical annotation performance can be found in Table S3 and S7.2. For the logistic regression classifier and the random forests classifier, this performance is nearly equal. For the linear SVM classifier, annotation performance sometimes drops for the non-greedy approach, especially for the ontology-annotated datasets. A possible explanation here could be that the linear SVM’s confidence scores are close together for the classes at the top of the hierarchy, which could result in different predictions for the greedy and non-greedy approach, as illustrated in Figure 1.C. This would imply that the confidence scores that are returned by the linear SVM classifiers are badly calibrated, which has been indicated in the literature before [Platt, 2000].

### 3.4 Technical variation can greatly affect flat and hierarchical performance without rejection, unlike biological variation

We also performed interdataset analyses with the help of the PbmcBench datasets (see Table S1) with flat and hierarchical annotation without rejection, to inspect the influence of biological and technical variation on the models’ performance. These analyses were performed following the same set-up as in Section 3.3 and the results can were visualised Figure S8.B. With interdataset analyses, the classifier is trained on one dataset and tested on an independent dataset. This setting resembles more real-life applications due to the presence of technical and biological variation. The PbmcBench datasets (see Table S1) contain two biological replicates that are sequenced with different protocols. Here, we trained the classifiers on the first biological replicate sequenced with 10Xv2 and tested on that replicate sequenced with different protocols and on the second biological replicate, which was also sequenced with 10Xv2.

The prediction results of the second biological replicate (pbmc2 10Xv2) show almost identical, high performance for both flat and hierarchical annotation, which indicates that biological variation does not influence the annotation process, as this result is the same as for the intradataset analyses. The results across the different protocols vary greatly, with sometimes flat annotation outperforming hierarchical annotation, sometimes the reverse, and sometimes equal performance. These observations align with those seen in a benchmark of automated annotation tools performed by Abdelaal et al. [2019] on the same dataset. In this benchmark, Abdelaal et al. [2019] concluded that interdataset annotation performance with technical variance depends on the protocol combination considered. So depending on the protocol combination, flat or hierarchical annotation can be the better choice.

### 3.5 Marker genes drive the annotation decisions of flat and hierarchical annotation

To complete the comparison of flat and hierarchical annotation without rejection, we also investigate whether or not they base their annotation decisions on biologically relevant features. We visualized the top-twenty (positive) coefficients of the flat model’s classifier and independent classifiers inside the non-greedy hierarchical annotation model for every label of the Azimuth PBMC dataset with the logistic regression classifier in Figures S10 - S17. Since the logistic regression classifier is a linear model, the coefficient scores it assigns to the features of the dataset, in this case the genes, reflect feature importance scores. These genes were then investigated with the help of CellMarker 2.0, a cell type marker gene database for several tissues in human and mouse [Hu et al., 2022]. Inspection of the AzimuthPBMC dataset was chosen as almost all the labels were present in the CellMarker 2.0 database and the logistic regression classifier was implemented, as it is the best performing classifier (see Fig. S8).

We find a substantial amount of marker genes associated with the top-twenty coefficients for flat annotation. This is also the case for hierarchical annotation, but the marker genes are spread out across the different independent classifiers. Important marker genes are identified at relevant parent nodes e.g. CD3A and CD3D at the T cell node and they are then absent at its (sub)children nodes. This indicates that the big annotation problem is correctly divided into biologically relevant sub-problems with the help of the hierarchy and that with this division, different division nuances are made in the different nodes. However, for some lower-level nodes no known marker genes are found back in the coefficients, while there were marker genes present for that label with flat annotation.

## 4 Discussion and conclusion

In this paper, we assessed the performance of three rejection settings in the context of cell type annotation for single-cell transcriptomics data: (1) no rejection, (2) partial rejection and (3) complete rejection, for both flat and hierarchical annotation. We observed considerable variation in rejection behavior across different classifiers and datasets throughout our analyses and so we strongly recommend inspection of the rejection behavior for a given annotation set-up to decide an appropriate rejection threshold. Accuracy-rejection curves prove to be an invaluable tool for this as exemplified in Figure 2. Our results also show that hierarchical annotation in comparison to flat annotation, leads to fewer label rejections under the full rejection setting, and these rejections are less severe under partial rejection. Consequently, if the rejection is implemented, hierarchical annotation proves to be the superior annotation method.

Furthermore, our study demonstrates that the implementation of partial rejection instead of full rejection substantially enhances the retained label information, irrespective of whether flat or hierarchical annotation methods are utilized. This demonstrates the usefulness of partial rejection in the context of single-cell annotation, as recently more and more extensive labeled singlecell atlases have become available with a wealth of knowledge that now can be optimally exploited with partial rejection [Elmentaite et al., 2022]. Moreover, as the bottom-up label assignment approach (i.e. partial rejection with flat annotation) is based on the output of normal ’flat’ annotation methods, this can be implemented after any existing ’flat’ annotation tool, marker-based or reference-based, that outputs reliable output scores together with the cell type annotation to further improve cell type annotation.

The comparison between flat and hierarchical annotation performance without rejection reveals very similar outcomes, with only technical variation contributing to substatial differences. Notably, in contrast to the other datasets, the performance of hierarchical annotation drops for the ontology-annotated Flybody and Flyhead datasets. These datasets are annotated with the FlyBase anatomical ontology (FBbt terms) [Gramates et al., 2022, Costa et al., 2013]. A possible explanation for this is that ontology-based hierarchies do not correctly represent transcriptomic relations between the cell type labels and are thus not useful for cell type annotation. This seems reasonable as the ontology hierarchies seem very complicated, with a lot of uninformative labels and labels that were confused by experts during manual annotation and are not located close to each other in the hierarchy. An interesting future field of research would be to explore several hierarchical structures for hierarchical annotation and construct cell type hierarchies based on single-cell transcriptomics data.

Lastly, we investigated whether or not flat and hierarchical annotation methods (without rejection) base their decisions on biologically relevant genes. We retrieved many marker genes that were pivotal for the decision process of flat and hierarchical annotation, but observed that marker genes were more spread out across the different classifiers for hierarchical annotation. For some labels at the bottom of the hierarchy, there were no marker genes found back in the top-twenty influential features of the lowest level classifiers for hierarchical annotation, while for these specific labels marker genes were found back with flat annotation. A possible explanation is that the differences that were picked up by the model did not necessarily correspond to marker genes but differentially expressed genes or genes with differential abundance. This can and does, as is proven in the performance of the hierarchical annotation, still lead to correct annotation but is not necessarily biologically intuitive for experts.

## 5 Data availability

The datasets were derived from sources in the public domain, they can be accessed through the repositories mentioned in the corresponding articles. Code to reproduce the analyses in this paper is freely available at: https://github.com/Latheuni/Hierarchical reject.

