## Supplementary Materials for "Uncertainty-aware single-cell annotation with a hierarchical reject option"

### 7 Supplementary Materials

#### 7.1 implementation details

Supplementary information

##### 7.1.1 Hyperparameters

Inside the flat and hierarchical annotation models, the LinearSVC(), the LogisticRegression() and the RandomForests() classifiers of scikit-learn [Pedregosa et al., 2011] were implemented. The hyperparameters implemented for these classifiers were the following:

- for the RandomForests() classifier:
  - n_estimators: 500
  - random_state: 0
  - max_depth: 30, 50 ,None
- for the LinearSVC classifier:
  - penalty: l2
  - C: 1, 10, 1000, 10000, 100000, 1000000
- for the LogisticRegression() classifier:
  - multi_class: multinomial
  - penalty: l2
  - max_iter: 2000

Highly variable gene (HVG) selection is performed with scanpy under the 'seurat' setting [Satija et al., 2015, Wolf et al., 2018]. For the F-test feature selection, the features were ranked according to their F-statistics and the number of selected features is considered as a hyperparameter. The tested hyperparameters range for the HVG selection was: [500, 1000, 2000, 5000, 8000, 1000] and for F selection: [1000, 5000, 10000, 15000, 18000, 20000], though sometimes one or several parameters were excluded due to running time problems. A detailed spreadsheet can be obtained through contact with the authors.

##### 7.1.2 Algorithms

The algorithms of greedy and non-greedy top-down label assignment and bottom-up label assignment are respectively defined in Algorithm S1, S2 and S3. Greedy and non-greedy label assignment (without rejection) corresponds to Algorithms S1, S2 when $t_r = 0$.

**Algorithm 1** Top-down greedy label assignment

**Input** test instance $\mathbf{x}$, a trained hierarchical model, rejection threshold $t_r$

$v \leftarrow \mathit{Root}$ , $P_{path} = 1$
**while** $v$ is not a leaf **do**
  $P_{opt}(\mathbf{x}) = 0$ , $v_{opt} \leftarrow v$
  **for** $v'$ in children($v$) **do**
    **if** $P(y \in v' | v, \mathbf{x}) > P_{opt}(\mathbf{x})$ **then**
      $P_{opt}(\mathbf{x}) \leftarrow P(y \in v' | v, \mathbf{x})$
      $v_{opt} \leftarrow v'$
    **end if**
  **end for**
  $v \leftarrow v_{opt}$
  $P_{path}(\mathbf{x}) \leftarrow P_{path}(\mathbf{x}) \times P_{opt}(\mathbf{x})$
  **if** $P_{path}(\mathbf{x}) < t_r$ **then**
    break
  **end if**
**end while**
**return** $v, P_{path}(\mathbf{x})$

**Algorithm 2** Top-down non-greedy label assignment

**Input** test instance $\mathbf{x}$, a trained hierarchical model, rejection threshold $t_r$

$Q \leftarrow$ Priority queue
$Q.addCouple(1.0, \mathit{Root})$
**while** $Q \neq \emptyset$ **do**
  $(P(\mathbf{x}), v) \leftarrow$ Q.popTopCouple()
  **if** $P(\mathbf{x}) < t_r$ or $v$ is a leaf **then**
    break
  **end if**
  **for** $v'$ in $children$(v) **do**
    $P'(\mathbf{x}) \leftarrow P(y \in v' | v, \mathbf{x})$
    $Q.addCouple(P(\mathbf{x}) \times P'(\mathbf{x}), v')$
  **end for**
**end while**
**return** $v, P(\mathbf{x})$

**Algorithm 3** Bottom-up label assignment

**Input** test instance $\mathbf{x}$, flat probabilistic classifier with associated prediction probability vector $P(y = c \,|\, \mathbf{x})$ for corresponding classes $\{c_1, ..., c_K\}$, rejection threshold $t_r$, a hierarchy $T$

$\mathcal{L} \leftarrow$ set that contains all leaves $\widehat{=}\{c_1, ..., c_K\}$
$v_{max} \leftarrow$ leaf that maximizes $P(y = c \,|\, \mathbf{x})$
$p_{max} \leftarrow$ cmax$(P(y = c \,|\, \mathbf{x}))$
**while** $p_{max} < t_r v_{max} \neq$ root **do**
    $\mathcal{L} \leftarrow$ all parents of nodes in $\mathcal{L}$
    **for** $v \in \mathcal{L}$ **do**
        compute $P(y \in v|\mathbf{x})$ using Eqn. (3)
    **end for**
    $v_{max} \leftarrow$ parent of $v_{max}$
    $p_{max} \leftarrow P(y \in v_{max}|\mathbf{x})$
**end while**
**return** $v_{max}, p_{max}$

##### 7.2 Supplementary tables

| Dataset | N° cells | N° genes | N° cell popu - lations | Protocol | Description | Balanced hierarchy | Label levels | Reference |
|---|---|---|---|---|---|---|---|---|
| AMB | 12 822 | 45 625 | 3/16/92* | SMART-seq v4 | Primary mouse visual cortex | Yes | 3 | Tasic et al. [2018] |
| COVID | 15 725 | 24 444 | 41 | NovaSeq v4 | BALF | No | 2-5 | Chan Zuckerberg Initiative Single-Cell COVID-19 Consortia et al. [2020] |
| Azimuth PBMC | 160 698 | 20 729 | 55** | CITE-seq | Human bone marrow | No | 2-5 | Stuart et al. [2019] |
| Flyhead | 53 809 | 13 056 | 71** | 10X | Head of the fruit fly | No | 3-9 | Li et al. [2022] |
| Flybody | 88 754 | 15 267 | 30** | 10X | Body of the fruit fly | No | 3-8 | Li et al. [2022] |
| PbmcBench: pbmc1_10Xv2 | 6444 | 33694 | 9** | 10X version 2 | PBMC | No | 1-3 | Ding et al. [2020] |
| PbmcBench: pbmc1_CL | 253 | 33694 | 7** | CEL-Seq2 | PBMC | No | 1-3 | Ding et al. [2020] |
| PbmcBench: pbmc1_DR | 3222 | 33694 | 9** | Drop-Seq | PBMC | No | 1-3 | Ding et al. [2020] |
| PbmcBench: pbmc1_iD | 3222 | 33694 | 7** | inDrop | PBMC | No | 1-3 | Ding et al. [2020] |
| PbmcBench: pbmc1_SM2 | 253 | 33694 | 6** | SMART-Seq2 | PBMC | No | 1-3 | Ding et al. [2020] |
| PbmcBench: pbmc1_SW | 3176 | 33694 | 7** | Seq-Well | PBMC | No | 1-3 | Ding et al. [2020] |
| PbmcBench: pbmc2_10Xv2 | 3362 | 33694 | 9** | 10Xv2 | PBMC | No | 1-3 | Ding et al. [2020] |



Table 1: Characteristics of the datasets used in this study after preprocessing (see Section 2.1).
* Number of cell type populations in every level of the hierarchy.
** Number of cell type populations at the lowest level of the hierarchy, for unbalanced datasets this lowest level is defined as the last sublabel available for every specific cell type population.

| | | AMB | COVID | Azimuth PBMC | Flyhead | Flybody |
|---|---|---|---|---|---|---|
| Greedy | LR | 0.927 | 0.806 | 0.910 | 0.917 | 0.956 |
| | RF | 0.853 | 0.686 | 0.798 | 0.832 | 0.928 |
| | SVM | 0.917 | 0.783 | 0.891 | 0.914 | 0.953 |
| Non-Greedy | LR | 0.927 | 0.806 | 0.904 | 0.919 | 0.955 |
| | RF | 0.841 | 0.689 | 0.799 | 0.832 | 0.928 |
| | SVM | 0.856 | 0.731 | 0.515 | 0.462 | 0.797 |

Table 2: The average accuracy scores across the five cross-validation folds of greedy and non greedy hierarchical annotation across the AMB, COVID, Azimuth PBMC, Flyhead and Flybody dataset for the logistic regression(LR), random forests(RF) and linear SVM classifier.

| | | AMB | COVID | Azimuth PBMC | Flyhead | Flybody |
|---|---|---|---|---|---|---|
| Greedy | LR | 0.835 | 0.594 | 0.803 | 0.831 | 0.829 |
| | RF | 0.660 | 0.329 | 0.481 | 0.628 | 0.562 |
| | SVM | 0.805 | 0.556 | 0.788 | 0.819 | 0.821 |
| Non-Greedy | LR | 0.834 | 0.594 | 0.788 | 0.829 | 0.823 |
| | RF | 0.662 | 0.343 | 0.488 | 0.628 | 0.562 |
| | SVM | 0.714 | 0.471 | 0.595 | 0.211 | 0.588 |

Table 3: The balanced accuracy scores of greedy and non greedy hierarchical annotation across the AMB, COVID, Azimuth PBMC, Flyhead and Flybody dataset for the logistic regression(LR), random forests(RF) and linear SVM classifier.

#### 7.3 Supplementary figures

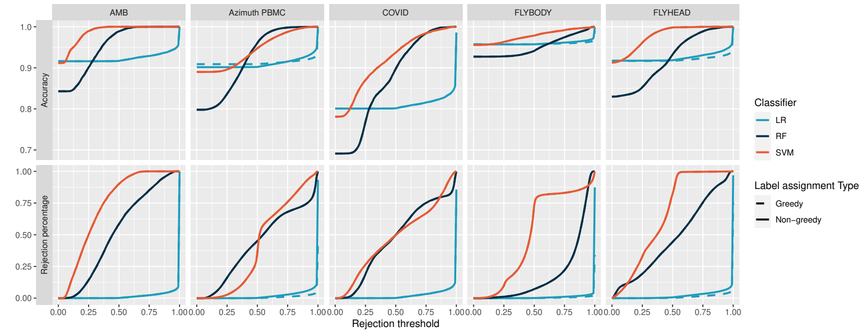


Figure 4: Accuracy-rejection curves of hierarchical annotation with greedy and non-greedy prediction of the AMB, Azimuth PBMC, COVID, Flybody and Flyhead datasets with Logistic regression (LR), Random forests (RF) and linear SVM (SVM) classifiers. The curves in the top row represent the accuracy score of the non-rejected label given a rejection threshold value. The bottom row curves represent the percentage of rejected labels for the rejection threshold value.Hierarchical rejection Greedy versus non-greedy.

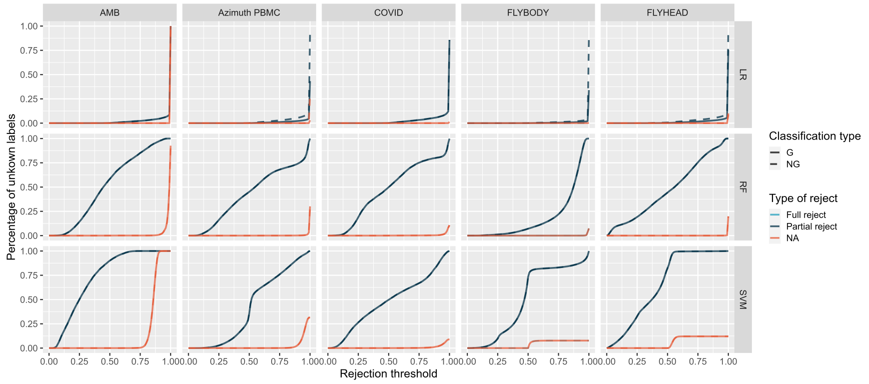


Figure 5: A comparison of the percentage of unknown labels assigned during the annotation task when partial and full rejection is implemented during greedy and non-greedy hierarchical annotation. Evaluation is performed with three classifiers (logistic regression, random forests and linear SVM) across 5 datasets (AMB, Azimuth PBMC, COVID, Flyhead and Flybody).

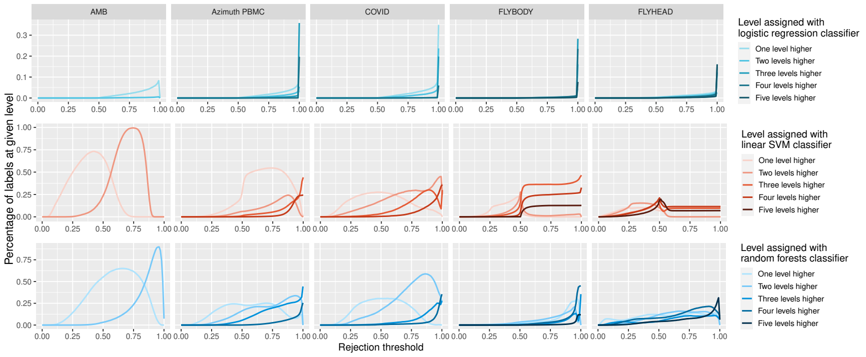


Figure 6: Details of partial rejection performed with non-greedy hierarchical annotation across a rejection threshold range from 0.0 to 1.0 for three classifiers (logistic regression - light blue, linear SVM - red, random forests - dark blue) across 5 datasets (AMB, Azimuth PBMC, COVID, Flyhead and Flybody). For all of these analyses, the percentage of labels that are assigned one level higher than the full label, two levels higher etc. is visualized.

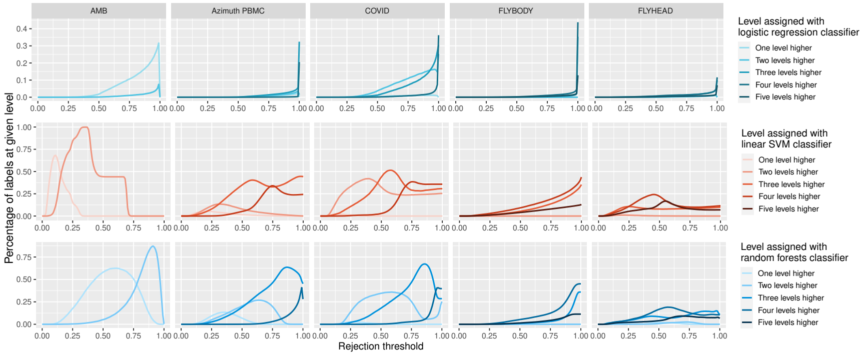


Figure 7: Details of partial rejection performed with flat annotation across a rejection threshold range from 0.0 to 1.0 for three classifiers (logistic regression - light blue, linear SVM - red, random forests - dark blue) across 5 datasets (AMB, Azimuth PBMC, COVID, Flyhead and Flybody). For all of these analyses, the percentage of labels that are assigned one level higher than the full label, two levels higher etc. is visualized.

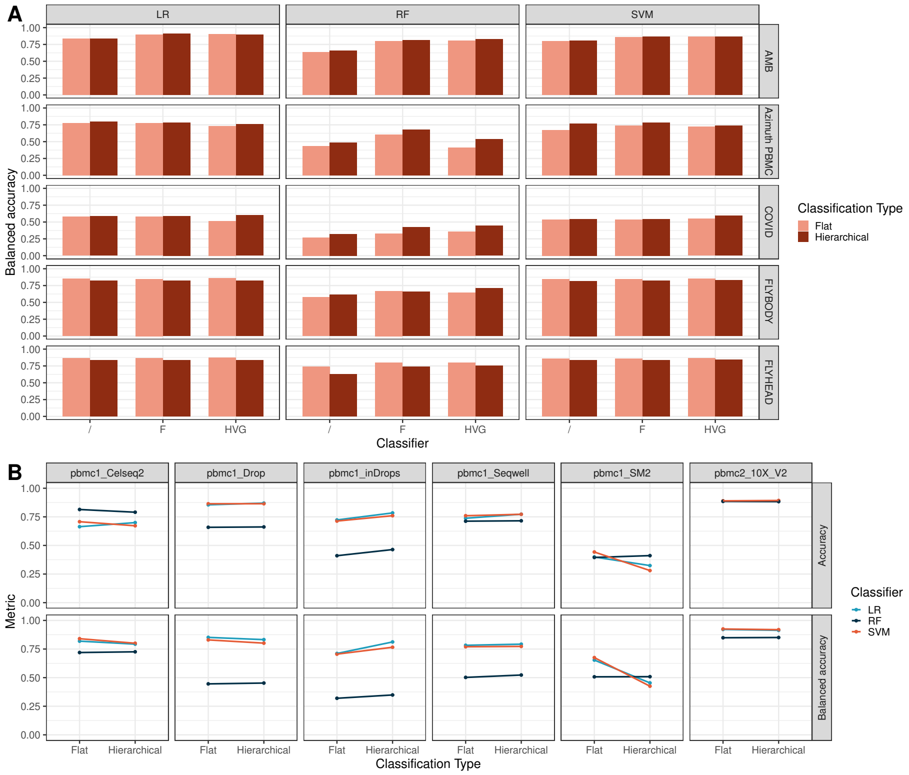


Figure 8: Flat versus hierarchical annotation performance without rejection on intra- and interdataset analyses. **A.** Comparison of the overall intradataset performance of flat versus hierarchical classification in terms of the balanced accuracy score for the AMB, AzimuthPBMC and COVID dataset (Table S1) with the use of logistic regression classifiers (LR), random forest classifiers (RF) and linear support vector machine classifiers (SVM). The classifiers are implemented with three feature selection strategies: no feature selection (/), the F-test (F) and highly variable gene selection (HVG). **B.** Performance of the hierarchical classifiers on intradataset analyses across every level in the hierarchy in terms of the balanced accuracy score across the different classifiers, datasets and feature selection methods. **C.** Interdataset performance of flat and hierarchical classification across the three classifiers without feature selection in terms of the accuracy and balanced accuracy score. The classifiers are trained on the pbmc1 sample, sequenced with the 10Xv2 technology. Performance is tested on sequenced data of the same sample across different plaforms and on a second biological sample (pbmc2), sequenced on the same platform.

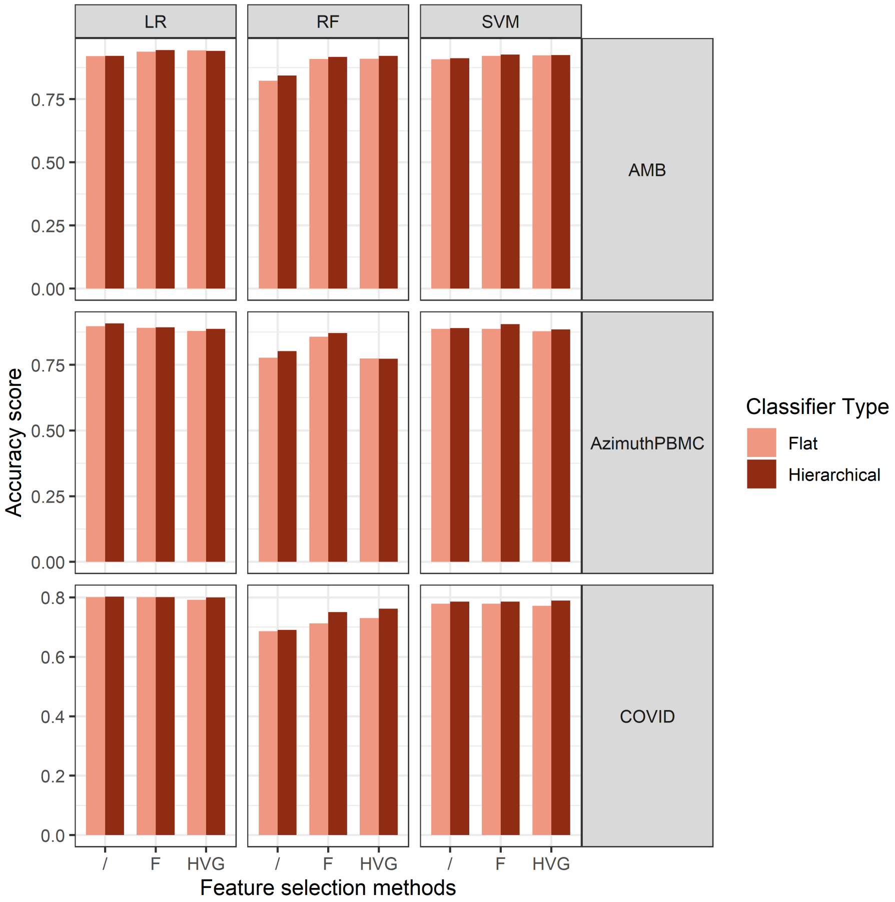


Figure 9: Comparison of the overall intradataset performance of flat versus hierarchical classification in terms of the accuracy score for the AMB, Azimuth-PBMC and COVID dataset (Table S1) with the use of logistic regression classifiers (LR), random forest classifiers (RF) and linear support vector machine classifiers (SVM). The classifiers are implemented with three feature selection strategies: no feature selection (/), the F-test (F) and highly variable gene selection (HVG).

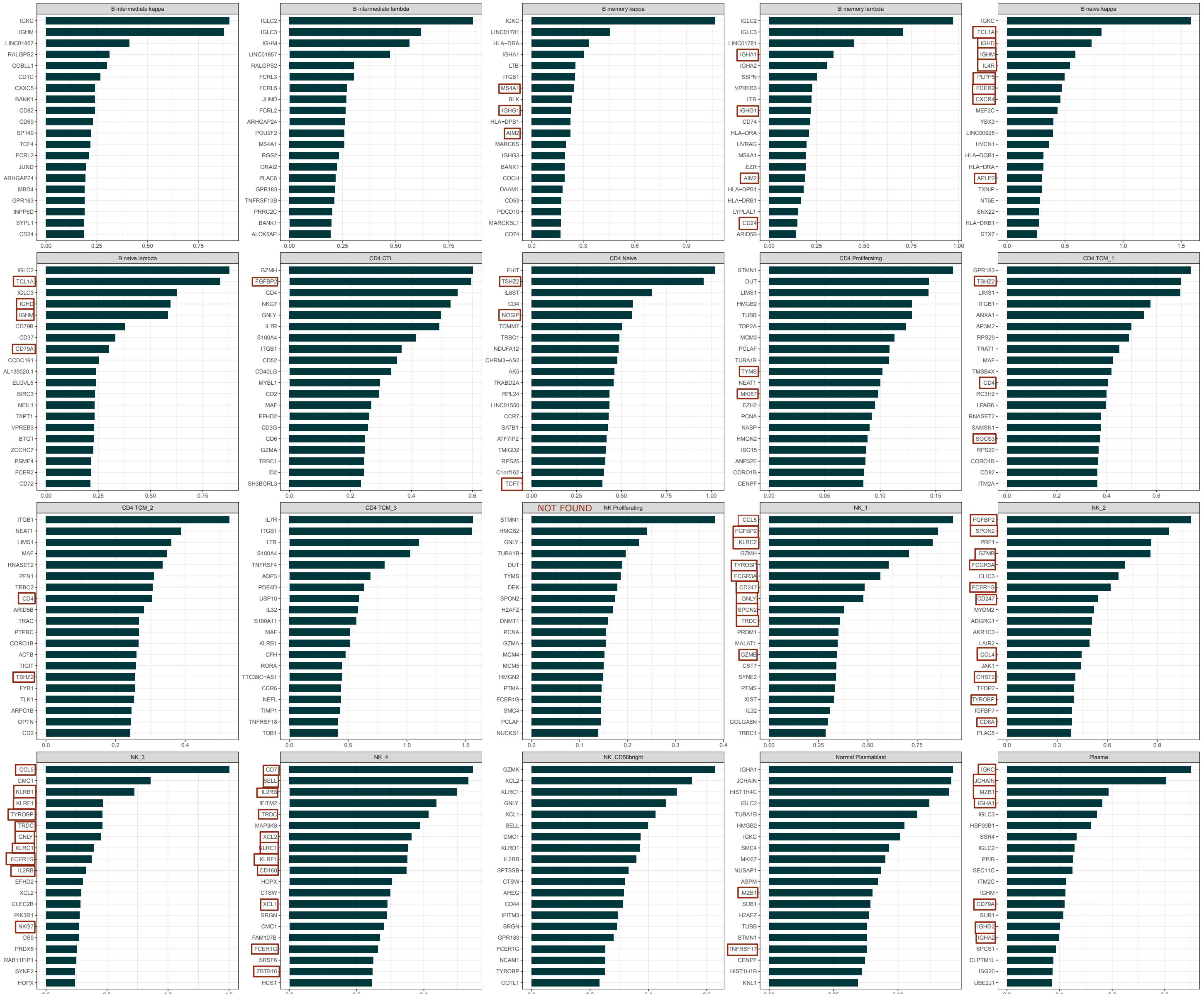


Figure 10: Part 1. Visualization of the top-twenty (positive) coefficients of flat annotation with logistic regression on the

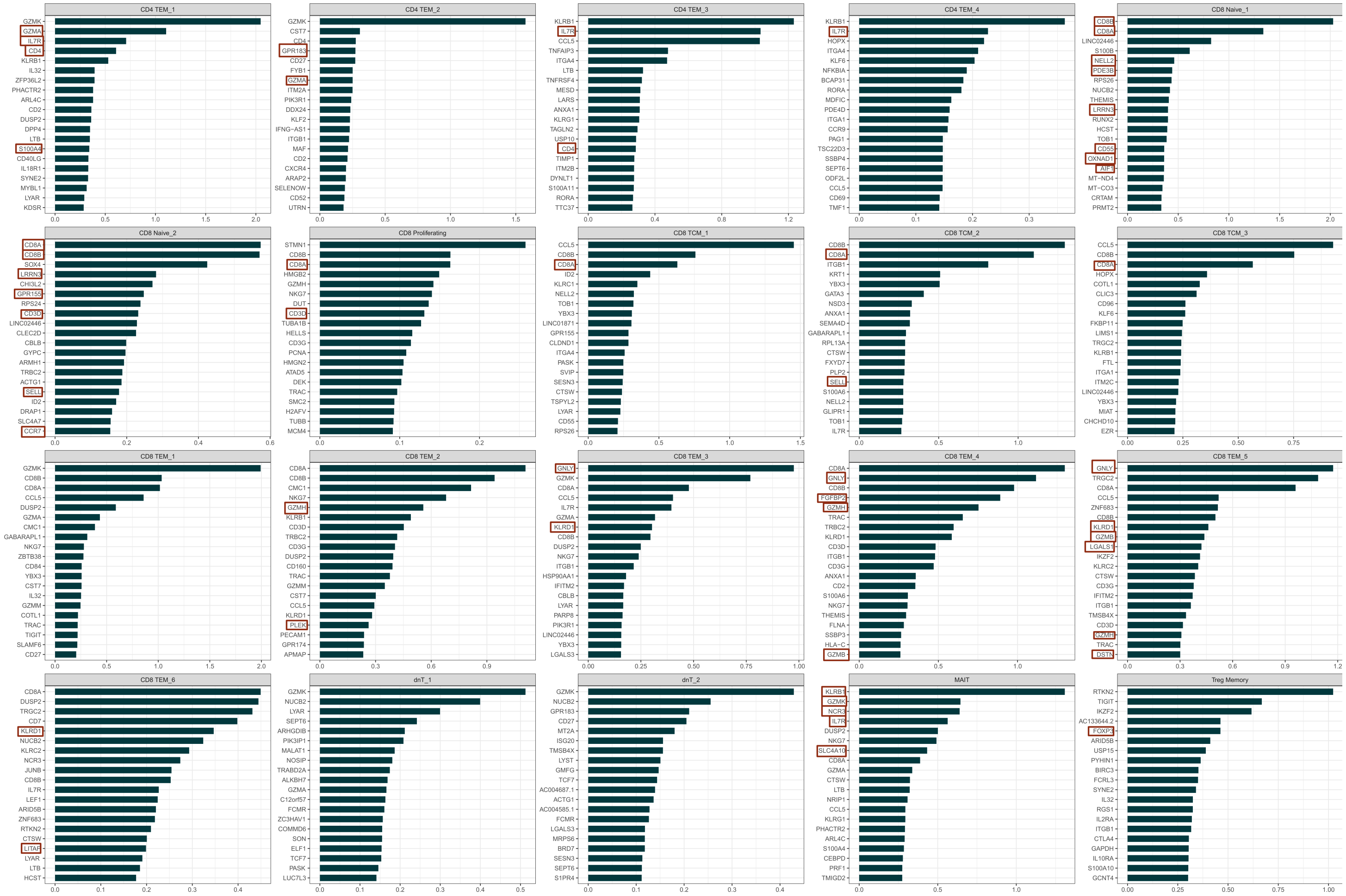


Figure 11: Part 2. Visualization of the top-twenty (positive) coefficients of flat annotation with logistic regression on the AzimuthPBMC dataset that leads to the final label classes. If the genes of these coefficients correspond to marker genes their names on the y-axis are enclosed by a red box.

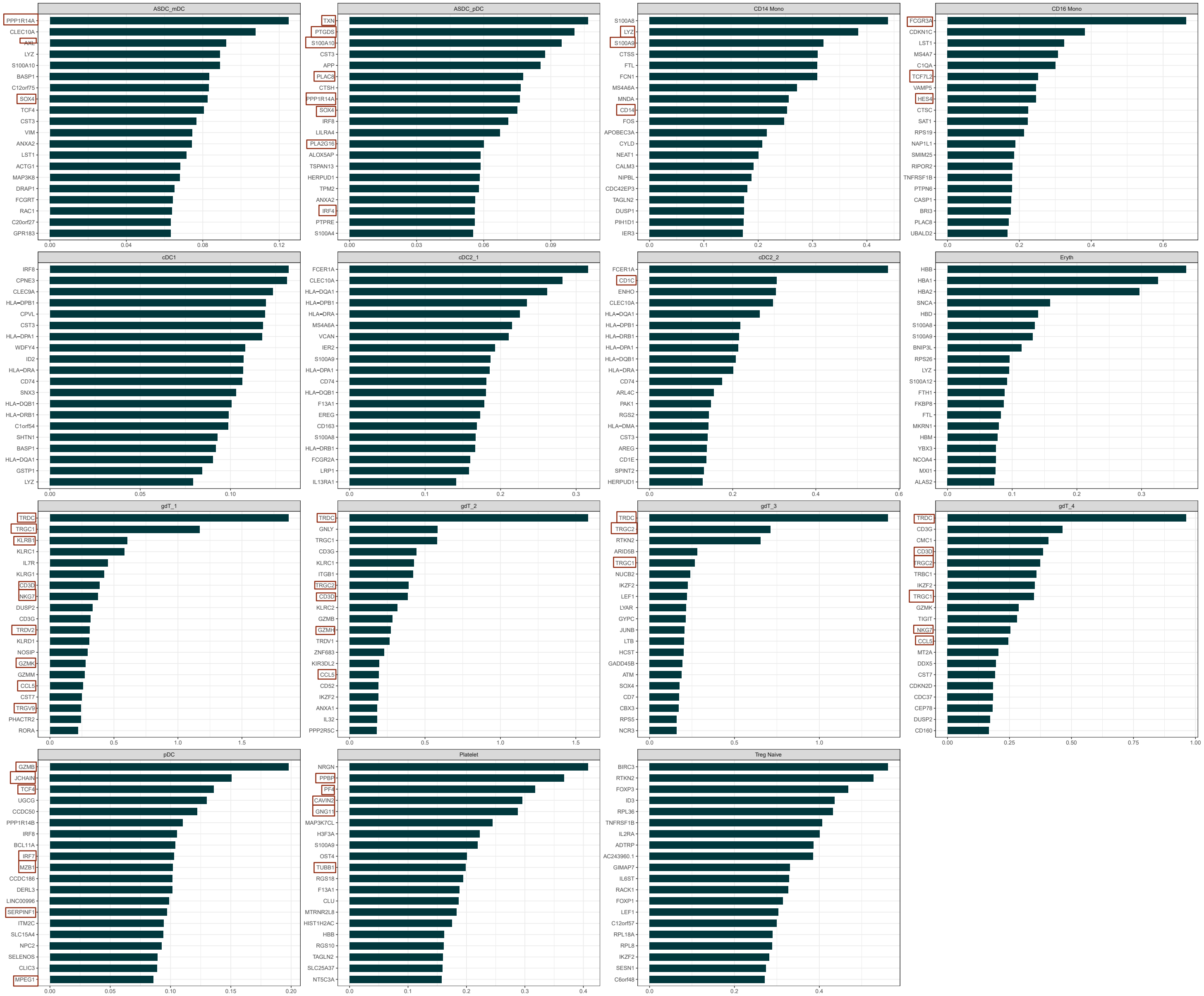


Figure 12: Part 2. Visualization of the top-twenty (positive) coefficients of flat annotation with logistic regression on the

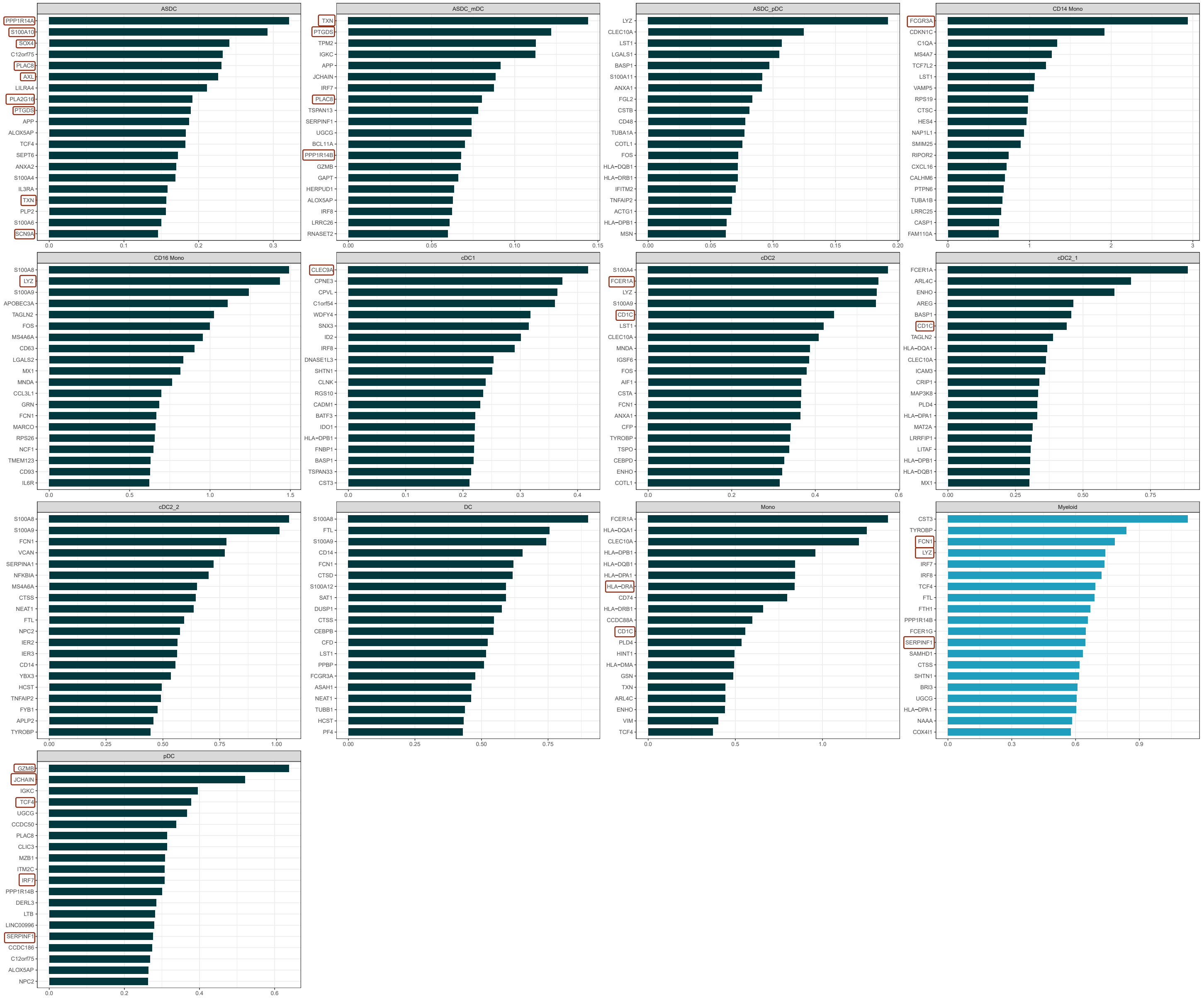


Figure 13: Visualization of the top-twenty (positive) coefficients of all myeloid sub- and super-labels for non-greedy hierarchical

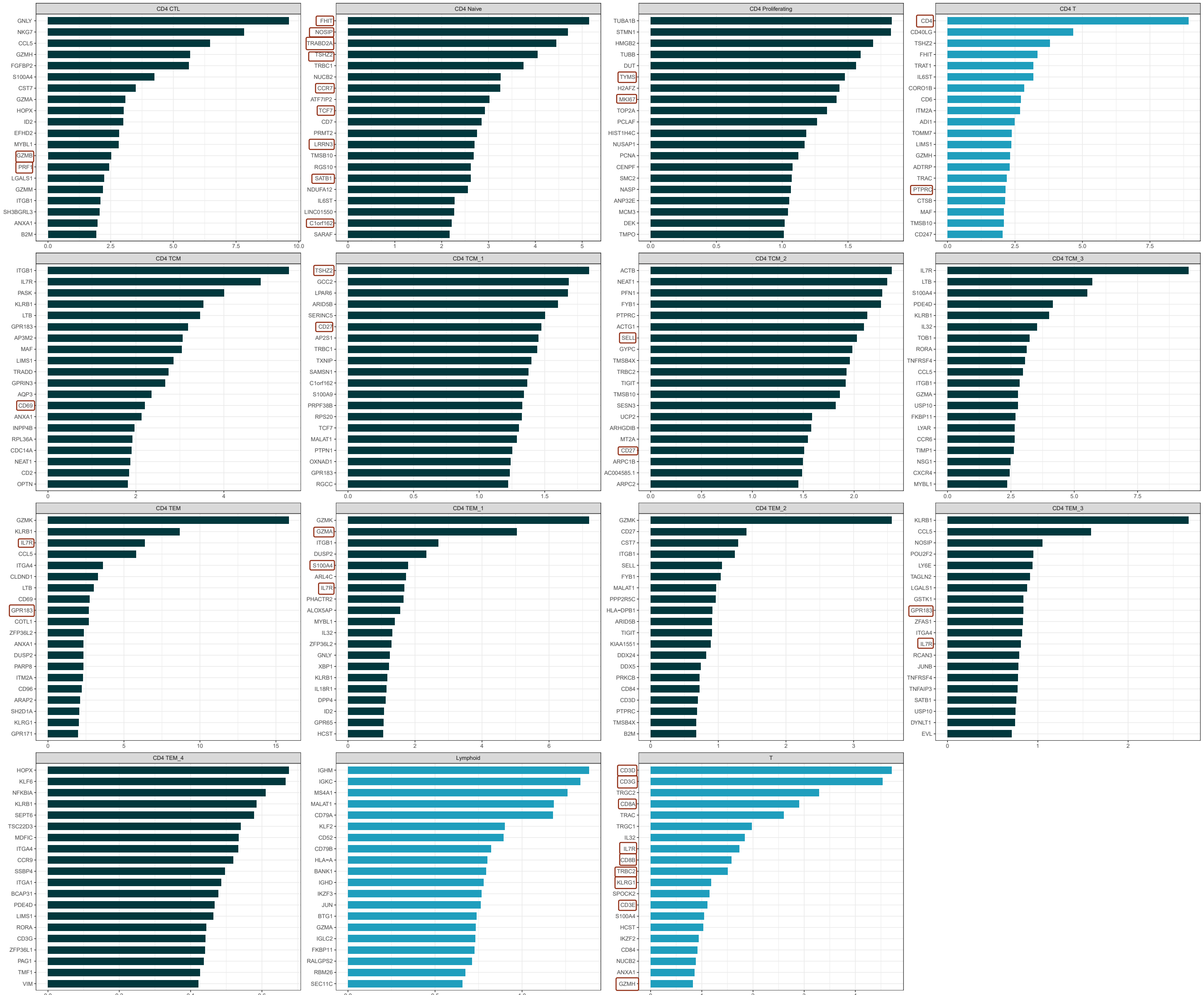


Figure 14: Visualization of the top-twenty (positive) coefficients of all CD4+ T cell sub- and super-labels for non-greedy

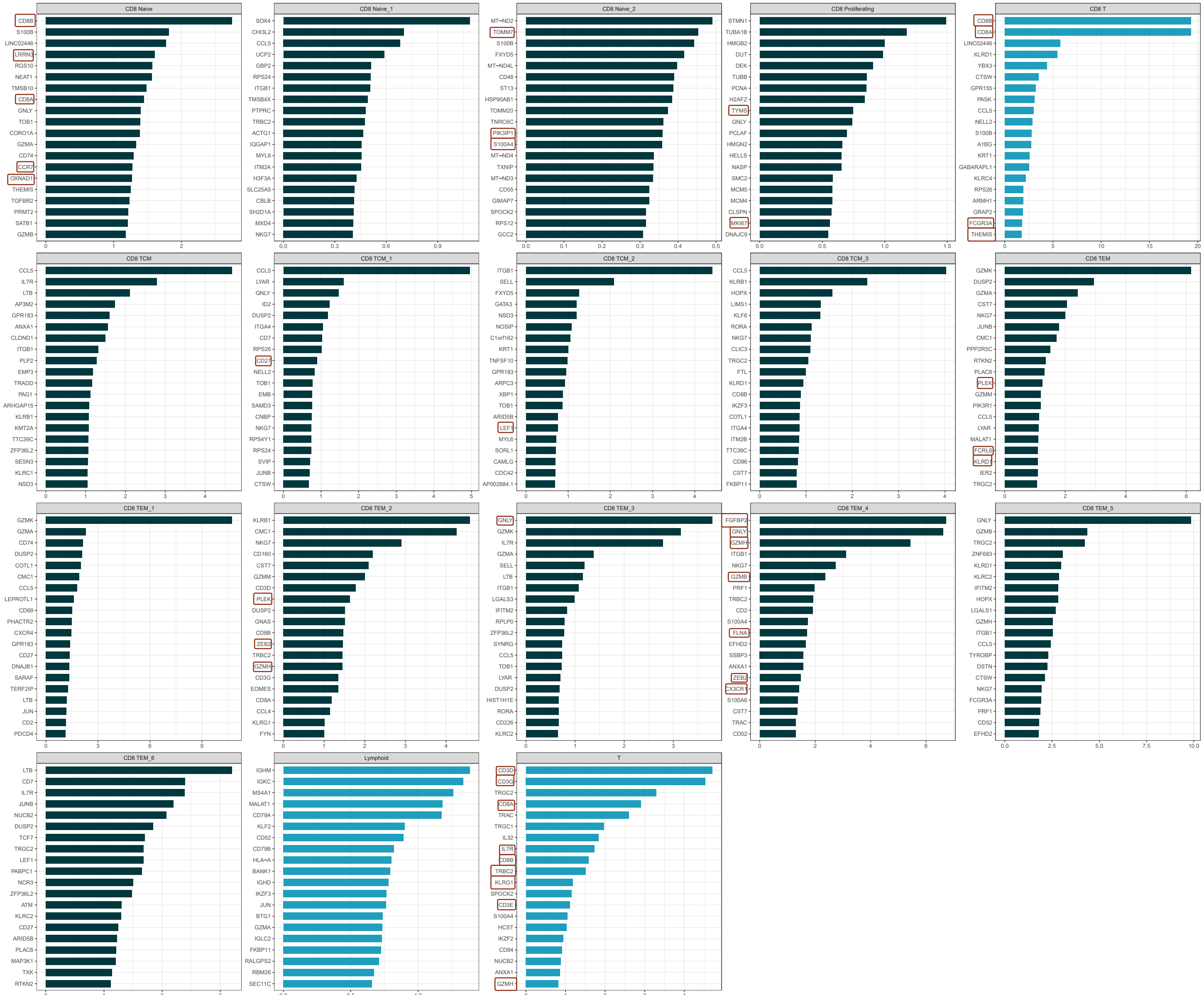


Figure 15: Visualization of the top-twenty (positive) coefficients of all CD8+ T cell sub- and super-labels for non-greedy

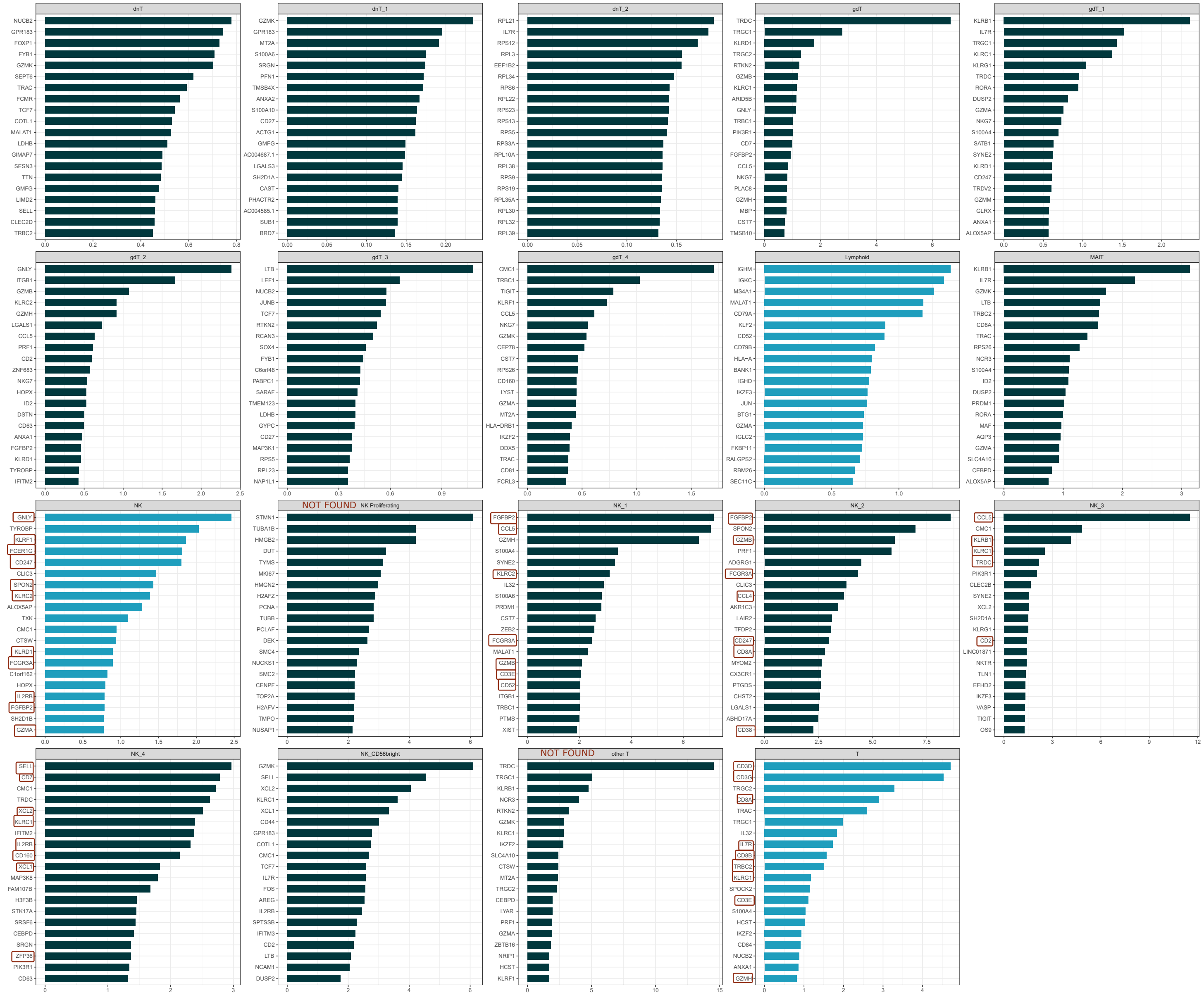


Figure 16: Visualization of the top-twenty (positive) coefficients of all other T and NK cell sub- and super-labels for non-greedy

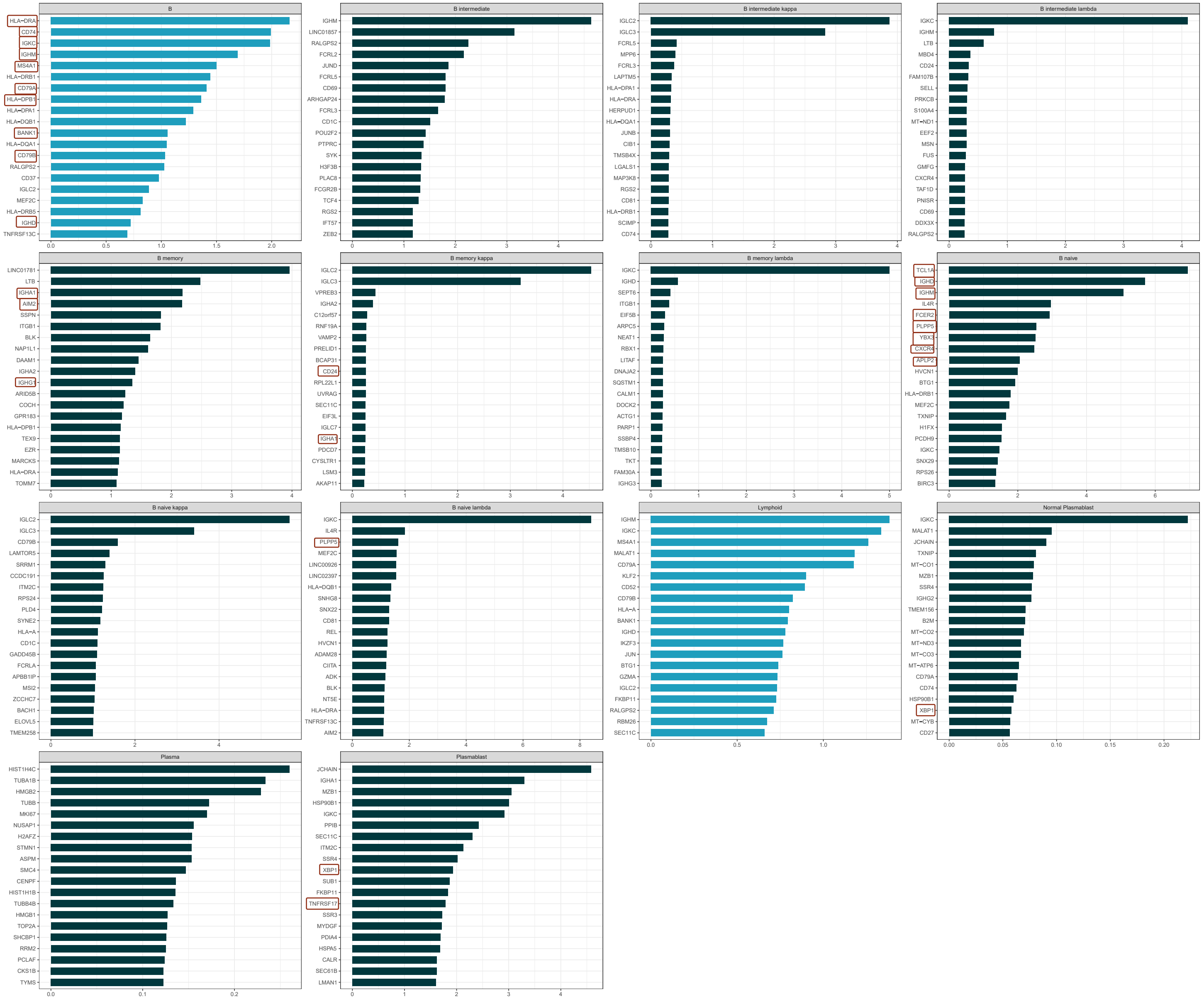


Figure 17: Visualization of the top-twenty (positive) coefficients of all other B cell sub- and super-labels for non-greedy

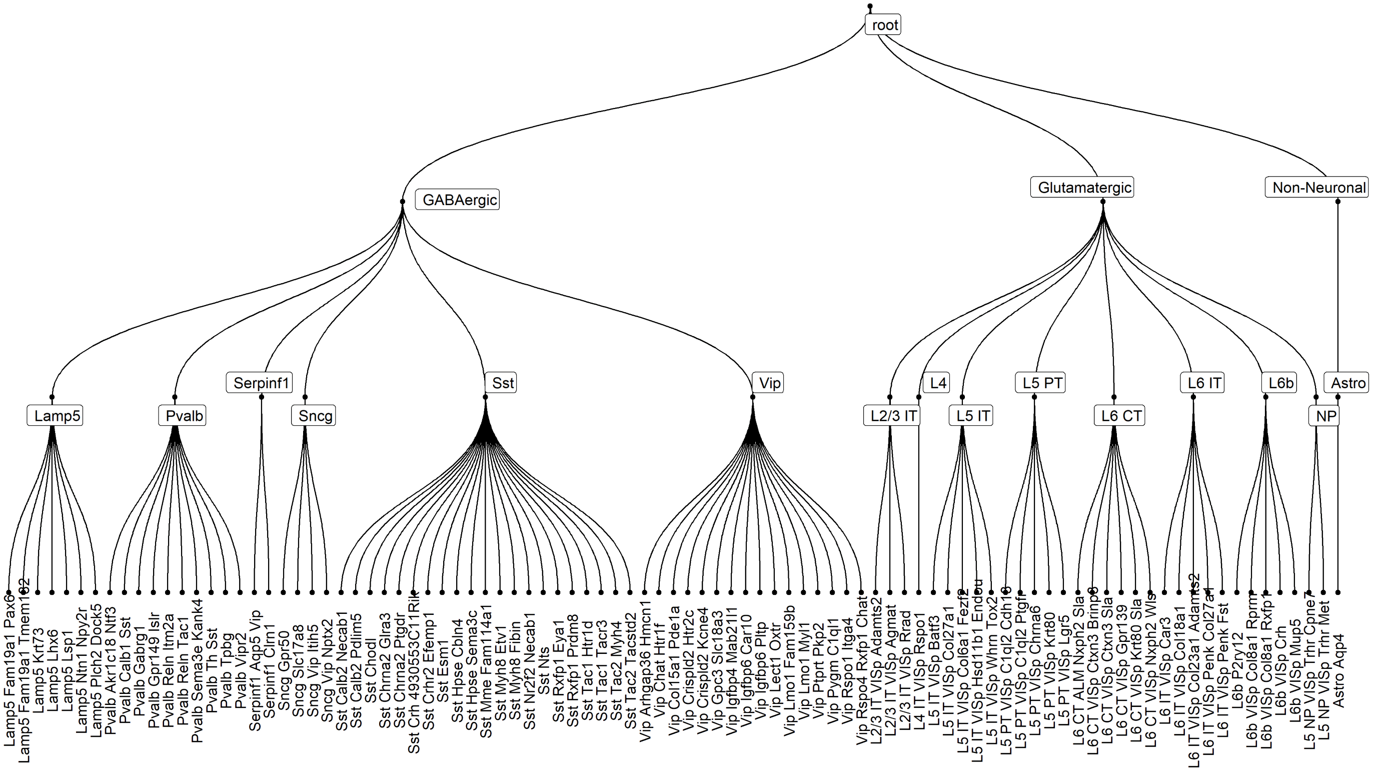


Figure 18: Visualisation of the hierarchy of the AMB dataset.

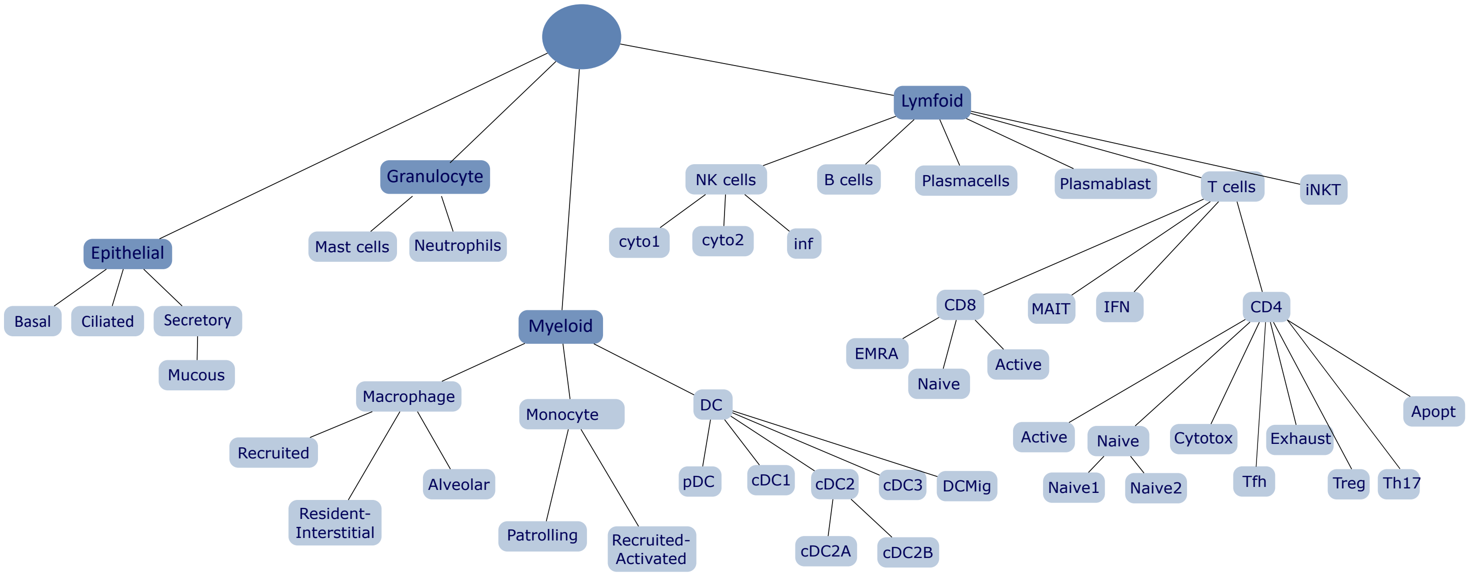


Figure 19: Visualisation of the hierarchy of the COVID dataset.

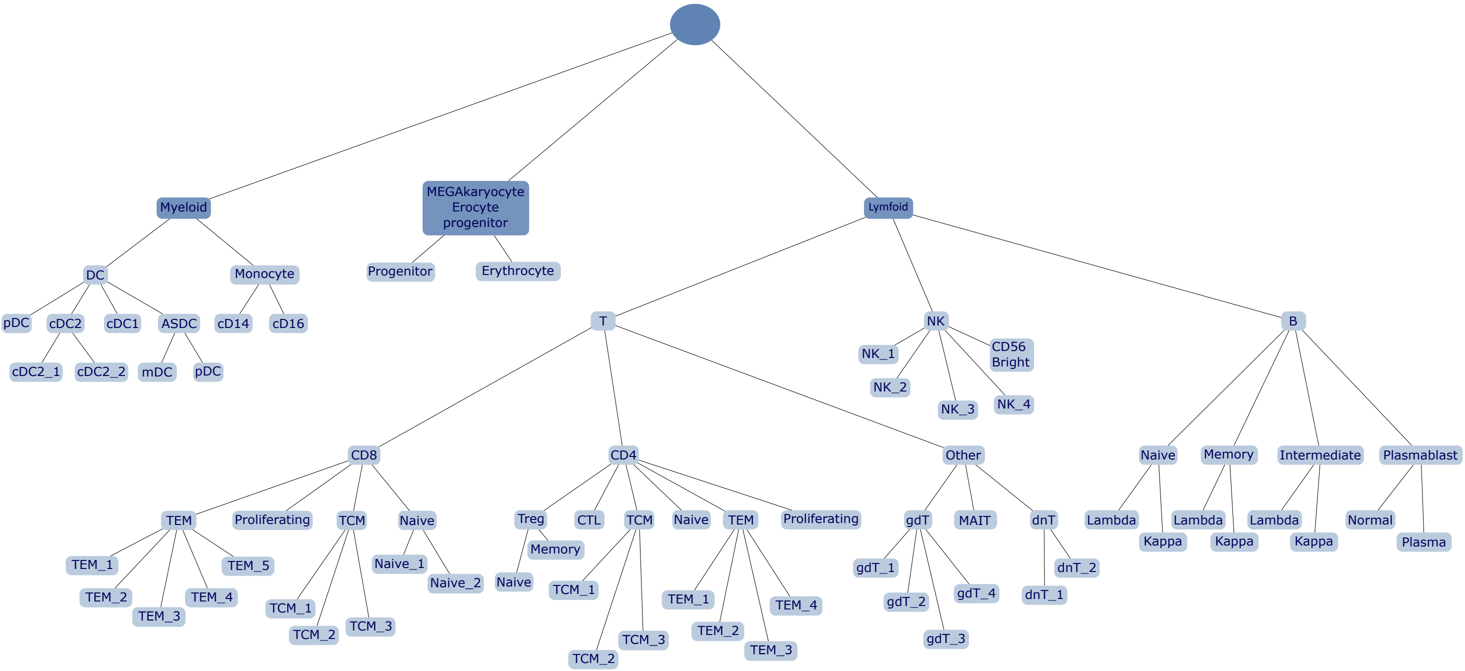


Figure 20: Visualisation of the hierarchy of theAzimuth PBMC dataset.

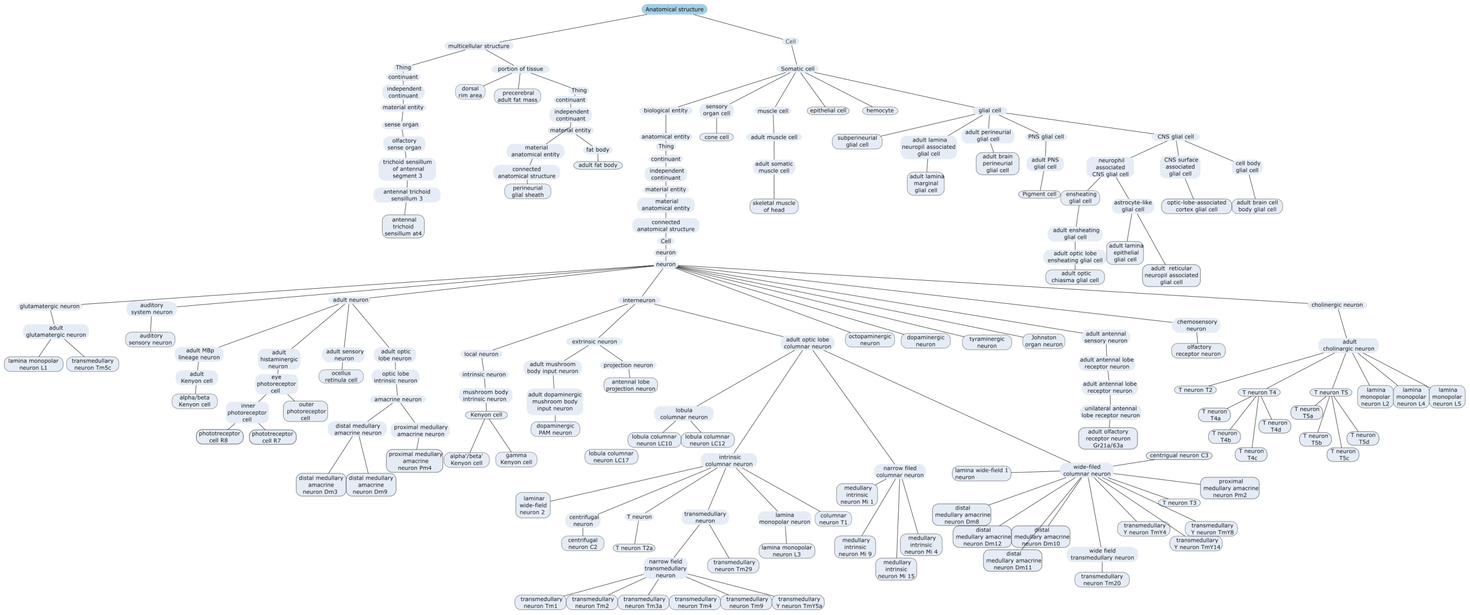


Figure 21: Visualisation of the hierarchy of theAzimuth PBMC dataset prior to exclusion of the uninformative labels.

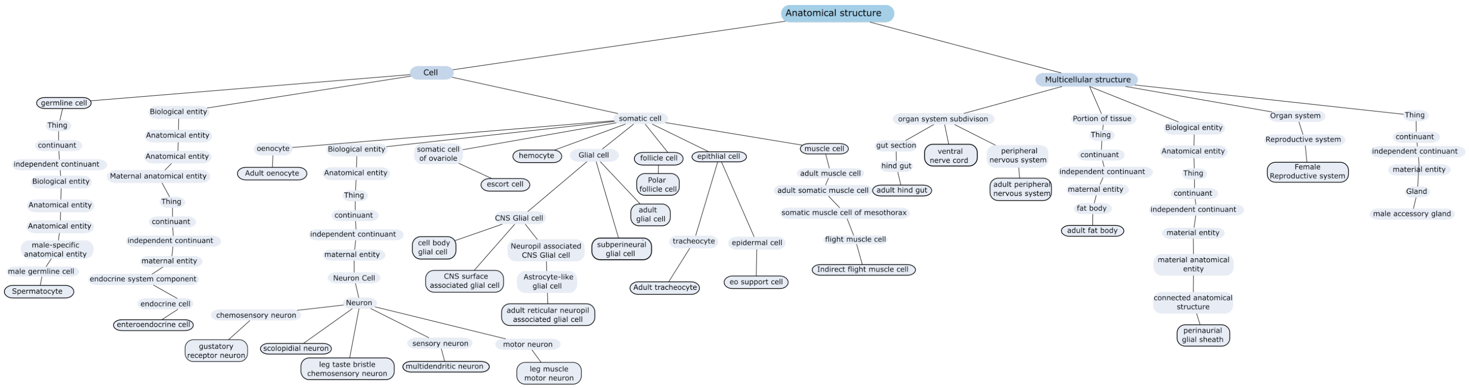


Figure 22: Visualisation of the hierarchy of theAzimuth PBMC dataset prior to exclusion of the uninformative labels.
